# Anxiety makes time pass quicker: neural correlates

**DOI:** 10.1101/2020.07.11.198390

**Authors:** I Sarigiannidis, K Kieslich, C Grillon, M Ernst, JP Roiser, OJ Robinson

**Affiliations:** UCL Institute of Cognitive Neuroscience; UCL Department of Clinical Education and Health Psychology; National Institutes of Mental Health, Bethesda, MD

## Abstract

Anxiety can be an adaptive process that promotes harm avoidance. It is accompanied by shifts in cognitive processing, but the precise nature of these changes and the neural mechanisms that underlie them are not fully understood. One theory is that anxiety impairs concurrent (non-harm related) cognitive processing by commandeering finite neurocognitive resources. For example, we have previously shown that anxiety reliably ‘speeds up time’, promoting temporal underestimation, possibly due to loss of temporal information. Whether this is due anxiety ‘overloading’ neurocognitive processing of time is unknown. We therefore set out to understand the neural correlates of this effect, examining whether anxiety and time processing overlap, particularly in regions of the cingulate cortex. Across two studies (an exploratory Study 1, N=13, followed by a pre-registered Study 2, N=29) we combined a well-established anxiety manipulation (threat of shock) with a temporal bisection task while participants were scanned using functional magnetic resonance imaging. Consistent with our previous work, time was perceived to pass more quickly under induced anxiety. Anxiety induction led to widespread activation in cingulate cortex, while the perception of longer intervals was associated with more circumscribed activation in a mid-cingulate area. Importantly, conjunction analysis identified convergence between anxiety and time processing in the insula and mid-cingulate cortex. These results provide tentative support for the hypothesis that anxiety impacts cognitive processing by overloading already-in-use neural resources. In particular, overloading mid-cingulate cortex capacity may drive emotion-related changes in temporal perception, consistent with the hypothesised role of this region in mediating cognitive affective and behavioural responses to anxiety.

## Introduction

Anxiety profoundly alters how we perceive the world (Grupe & Nitschke, 2013; Robinson et al., 2013), promoting harm-avoidant behaviours. Anxious individuals tend to pay more attention to threatening stimuli (e.g. attentional bias: Cisler & Koster, 2010; Robinson et al., 2014; Van Bockstaele et al., 2014), interpret ambiguous information as threatening (e.g. interpretation bias: Wilson et al., (2006)) and overestimate the probability and personal cost of negative events (e.g. judgement bias: Aylward et al., 2019; Charpentier et al., 2017; Mitte, 2007). While previous behavioural and neuroimaging work has mainly focused on how anxiety influences emotional information (i.e. hot cognition) (Bar-Haim et al., 2007; Carlisi & Robinson, 2018; Cisler & Koster, 2010; Mathews et al., 1997), less research has been conducted on how anxiety influences non-emotional information (i.e. cold cognition) with mixed results (Robinson et al., 2013). To this end, in a series of prior studies we demonstrated that anxiety (induced in healthy individuals using threat of unpredictable shock) reliably leads to alterations in non-emotional perception, namely temporal perception (Sarigiannidis, Grillon, et al., 2020). We found clear evidence that anxiety leads to underestimation of time, i.e. that time speeds up under threat of unpredictable electric shock, possibly due to the loss of temporal information (Sarigiannidis, Grillon, et al., 2020; Sarigiannidis, Kirk, et al., 2020). In the present study we extend this work to probe the neural correlates of the influence of anxiety on temporal processing using a similar bisection task.

Previous functional magnetic resonance imaging (fMRI) studies employing a similar anxiety manipulations to ours (Kirlic et al., 2017, 2017; McMenamin et al., 2014) have consistently found activation in the anterior insula while participants passively anticipate unpredictable shocks (see also: Robinson et al., 2019). Other brain areas that have been shown to be activated during sustained threat include the anterior and posterior cingulate gyrus, thalamus, caudate and cerebellum (Mechias et al., 2010; Robinson et al., 2019). These brain areas have also been associated with processing and anticipating painful stimuli (Koyama et al., 2005; Wager et al., 2013), supporting the hypothesis that they may play a general role in prolonged states of negative anticipation. Consequently, we would expect to replicate these activations in the present study when individuals are under threat of shock-induced anxiety.

A broad network of brain regions has been reported to be recruited during time perception. A recent study suggested that similarly to sensory cortical maps, topographic timing maps exist; where different brain areas respond to specific ranges of temporal intervals, and whose selectivity changes gradually (Harvey et al., 2020). For supra-second intervals (the focus of the current work), cortical brain regions are more heavily involved (Nani et al., 2019; Wiener et al., 2010), including the cingulate and frontal cortex, as well as the pre-supplementary motor area (pre-SMA) which is considered central (Schwartze et al., 2012).

There have been no studies exploring the interaction between anxiety and time perception at the neural level. However, more broadly, previous studies have highlighted the involvement of frontal areas in the interaction between anxiety and tasks tapping into cold cognition (Bishop, 2007, 2009; Carlisi & Robinson, 2018; Robinson et al., 2019). Consistent with this, two prior threat-of-shock studies found that anxiety increased activation in frontal areas (including the superior frontal gyrus), and these activations were also associated with anxiety-related behavioural change (Balderston et al., 2017; Torrisi et al., 2016). However, assuming that the effect of anxiety on cognitive function can be likened to classic multitasking interference (where two tasks compete for limited cognitive and neural resources, and hence interfere with one another: Eysenck et al., 2007; Watanabe & Funahashi, 2018), we would not necessarily anticipate that there exists a specific brain region underlying interference between anxiety and cognition. Instead, the precise region(s) implicated may depend on the particular cognitive task used to probe anxiety, since multitasking interference is a result of overlapping resources (Maillet et al., 2019; Nijboer et al., 2014; Watanabe & Funahashi, 2018). We have previously argued that the impact of anxiety on time perception is driven by demands on attention, a cognitive resource both of these processes might be utilising (Sarigiannidis, Grillon, et al., 2020; Sarigiannidis, Kirk, et al., 2020). Thus, it is possible that the impact of induced anxiety on temporal perception is driven by overlap between time- and anxiety-related neural processing. A strong candidate brain area for such overlap is the pre-SMA. Time-perception related activations in the pre-SMA have been reported to vary parametrically with the amount of attention allocated to timing a stimulus (Coull et al., 2004). At the same time, the pre-SMA seems to be activated by threat-of-shock manipulations as revealed by a meta-analysis (Chavanne-Arod & Robinson, 2020).

To test our hypothesis, we initially conducted a small exploratory study (Study 1), which was used to refine our design and generate pre-registered (Sarigiannidis, 2019) predictions for the second study (Study 2). Importantly, for Study 2, we calibrated our temporal cognition task to each participant to exclude the possibility that any neural differences observed were due to the properties of the different temporal intervals of the task. Our specific predictions for Study 2, were the following:

1. Induced anxiety would lead to temporal underestimation, replicating our previous finding (Sarigiannidis, Grillon, et al., 2020). Specifically, we predicted that participants would perceive the temporal intervals as shorter when under threat of shock.
2. Anxiety would elicit activation in the anterior cingulate cortex and the caudate (in Study 2), as identified by our pilot study (Study 1).
3. Time perception would elicit activation in the pre-SMA and right inferior frontal gyrus, as identified in a previous meta-analysis (Wiener et al. (2010).
4. Time-related and anxiety-related neural processing would interact in the pre-SMA
5. Threat-induced activation changes during temporal processing would correlate with the threat-induced changes in time estimation.

## Procedure

### Overview

All studies consisted of a single testing session. Following written informed consent, as approved by local ethical procedures (see below for specifics), and the completion of questionnaires, a shock calibration procedure was completed by the participant in the scanning room to determine an appropriate level of aversive electrical stimulation. Participants then completed the temporal bisection task under threat of unpredictable shock and safe conditions inside the scanner. During each one-hour scan, anatomical and functional images were acquired, with each of the two functional runs lasting approximately 15 minutes. During the task, participants selected between two responses (short and long) in a two-alternative forced choice manner inside the scanner, via an MRI-compatible button box. Information relating to participant recruitment and inclusion/exclusion criteria is provided in each of the experiment-specific methods sections below.

### Study Site

Study 1 was completed at the National Institutes of Health, Bethesda, MD, USA, while Study 2 was completed at University College London (UCL), UK.

### Apparatus

In Study 1, experiment material was presented on Windows computers using E-prime, while Study 2 was run in Cogent 2000 (www.vislab.ucl.ac.uk/cogent.php; Wellcome Trust Centre for Neuroimaging and Institute of Cognitive Neuroscience, UCL, London), under Matlab.

### Shock calibration

A shock calibration procedure was performed prior to testing in order to control for shock tolerance and skin resistance. Single pulse shocks (Study 1) or trains of shocks (Study 2) were delivered to the non-dominant wrist via a pair of silver chloride electrodes using a DS7 (Study 1) or a DS5 (Study 2) stimulator (Digitimer Ltd, Welwyn Garden City, UK). Participants received shocks sequentially with step increases in amplitude, which they rated using a scale from 1 to 10 (1 labelled “I barely felt it” and 10 labelled “approaching unbearable”). The level of shock delivered in the experiment was set to 80% of the maximum tolerated for each individual.

### Temporal bisection task under threat of shock: Overview

In both Study 1 and 2, participants completed a visual temporal bisection task under two alternating conditions (Figure 1): “threat-of-shock” (labelled “threat”), during which they could receive shocks at any time without warning, and “safe” during which they could not receive any shocks (the order was counterbalanced). The task was flanked by coloured borders that indicated the condition (safe or threat), taken from a pool of four colours (red, blue, green, magenta), which was also counterbalanced across participants.

**Figure 1:**
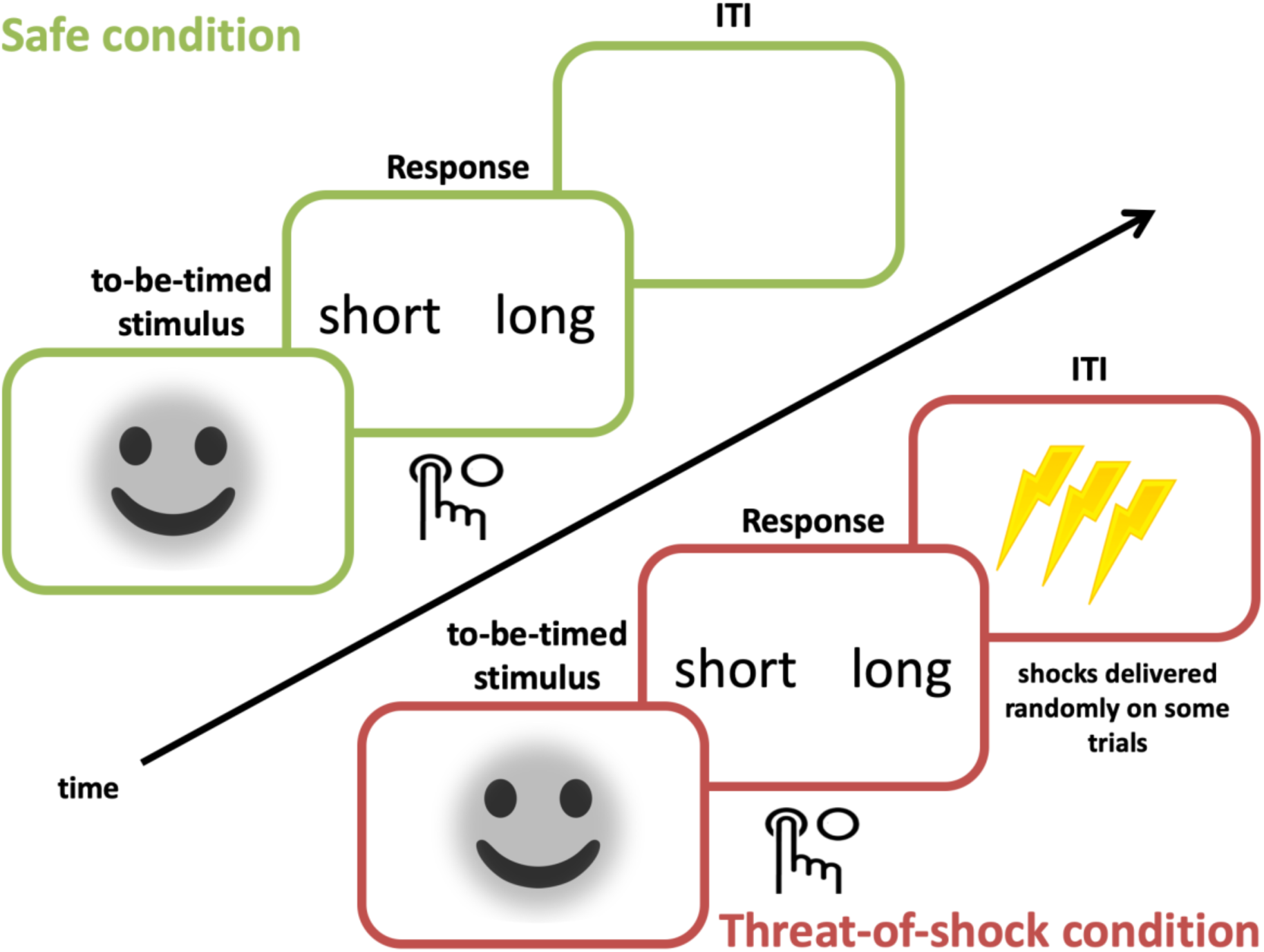
Experimental parameters for the studies and demographic information. Stimulus durations differed between Study 1 and 2 (see below, Experiment specific methods). Participants were given 1.5s to respond and the inter-trial intervals (ITIs) were 0.5, 1.1, 1.7, 2.3, 2.9, 3.5s. Note: in the actual experiment participants were presented with NimStim images (Tottenham et al., 2009).

A short training phrase preceded the main task. It consisted of presenting participants with two anchor durations (Figure 1), a “short” duration (1.4s) and a “long” duration (2.6s). Each was shown three times, and presentation order was pseudorandomised. In addition, before the beginning of each block (safe or shock) the anchor durations were repeated to ensure consolidation.

The to-be-timed stimuli were pictures of emotional facial expressions (happy, fearful or neutral; taken from a standardised set: Tottenham et al., 2009). Stimulus durations differed between Study 1 and 2 (see below, Experiment specific methods). All stimulus durations were pseudorandomised, and presented equally often in each threat and safe block to avoid potential biases (Wearden & Ferrara, 1996). On each trial participants were required to: press “short” if the duration of the stimulus was more similar to the “short” anchor, or press “long” if the duration of the stimulus was more similar to the “long” anchor (left and right buttons for these options were counterbalanced across participants). After the 1.5s response limit, there was a variable inter-trial interval (ITI: 0.5, 1.1, 1.7, 2.3, 2.9, 3.5s, pseudo-randomised). Participants were explicitly told to avoid counting seconds as well as avoid any other strategy to estimate the duration of the stimuli; instead they were asked to make the temporal judgments based on their gut feeling.

### Psychophysical modelling

We fitted psychometric functions to each participant’s data, separately for the safe and the threat condition in order to calculate the bisection point (BP). The BP is defined as the stimulus duration that is perceived by the participant to be of equal distance to the short and the long anchor, i.e. the stimulus duration that corresponds to 50% of the participants proportion of long responses, pLong (Kingdom & Prins, 2010). The BP is used as a measure of the perceived duration of comparison intervals. For example, a rightward shift of the psychophysical curve would lead to a greater BP, indicating underestimation of time (and vice-versa for a leftward shift).

The data was modelled using the Palamedes toolbox in Matlab (Prins & Kingdom, 2009) using maximum likelihood estimation. The proportion of long responses, pLong, at each comparison interval, was fitted with logistic functions defined by four parameters: threshold α, slope β, guess rate γ, and lapse rate λ. In line with previous studies, γ was fixed at 0 since the task was 2-alternative forced-choice; λ was fixed at 0.1 to allow for occasional attentional lapses (Terhune et al., 2016); and α and β were free parameters. The duration corresponding to the 50% threshold on the psychometric function was defined as the BP.

## Experiment specific methods

### Study 1: Temporal bisection task under threat of shock

The fMRI study and all procedures were approved by the NIH Institutional Review Board Project (ID Number: 02-M-0321) and were conducted in accordance with the Declaration of Helsinki. The aim of Study 1 was to identify regions of interest (ROIs); 13 individuals were scanned.

Participants were recruited through advertisements (newspaper and public transport) in the Washington, D.C. metropolitan area. Following an initial telephone screen, individuals visited the NIH for comprehensive screening by a clinician, which comprised a physical examination, urine drug screen, and the Structured Clinical Interview for the Diagnostic and Statistical Manual of Mental Disorders, Fifth Edition (American Psychiatric Association et al., 2013). Exclusion criteria were: contraindicated medical disorder (i.e. those thought to interfere with brain function and/or behaviour); past or current psychiatric disorders; and use of psychoactive medications or recreational drugs (per urine screen). All participants provided written informed consent and were reimbursed for their participation.

In Study 1, the duration of the to-be-timed stimuli varied between 1.4-2.6s (six possible durations: 1.4, 1.64, 1.88, 2.12, 2.36, 2.6s). The session consisted of two runs, each comprising of four blocks, two in the safe and two in the threat condition (counterbalanced) with 36 trials per block.

Seventy-two pictures were used in this experiment, depicting happy, neutral and fearful facial expressions, taken from 24 actors. During each block, participants viewed an equal number of happy, fearful and neutral facial expressions, the order of which was pseudorandomised. Similarly, stimulus durations were pseudorandomised within each block, so that all durations were repeated six times.

Participants received between 0-3 shocks during each threat block, only during threat blocks. The order of the shocks was random for each participant and occurred on different trials, following the participants’ response.

Scanning was performed on a 3T Siemens Magnetom Skyra using a 32-channel head coil. Each volume consisted of 34 slices and was acquired in TR=2s. Other parameters of the functional EPI were the following: voxel size of 3mm × 3mm (slice thickness of 2.5mm, 0.5mm slice separation), field of view of 216mm × 216mm, echo time of 30ms; flip angle of 70°.

### Study 2: Modified temporal bisection task under threat of shock

The studies and all procedures were approved by the UCL Research Ethics Committee (Project ID Number: 1227/001) and were conducted in accordance with the Declaration of Helsinki. Prior to the fMRI study we completed a behavioural pilot to ensure that the properties of the modified task were as expected. We pre-registered this study (Sarigiannidis, 2019).

Participants were recruited from UCL databases. A telephone interview was conducted to screen for past neurological or psychiatric diagnosis and to determine MRI safety. In addition, on the day of testing participants completed self-report measures of depression (Beck Depression Inventory: BDI; Beck & Steer (1987)) and trait anxiety (State Trait Anxiety Inventory: STAI; Spielberger, 1983) to ensure they fell within the non-clinical range. Participants provided written informed consent and received £20 for the fMRI study (which lasted approximately two hours: one hour for task explanation and questionnaires, and one hour for scanning).

A power calculation (G*power version 3.1.9.2 (Faul et al., 2007)) determined the sample size based on the meta-analytic effect of threat (d=0.68) from our previous behavioural temporal bisection studies (Sarigiannidis, Grillon, et al., 2020). Given that the MRI scanner may be an anxiogenic environment, which could raise anxiety levels during the safe condition and therefore reduce condition differences, we decreased the expected effect size by ∼30% (d=0.47). In order to achieve d=0.47 with 80% power and an alpha of 0.05 (two-tailed), the required sample size required was 30 participants. In the fMRI study, one participant was excluded from analysis as they pressed the same button on every trial, leaving us with a final sample size of 29.

Before the main task, participants performed the calibration task whose trial structure was identical to that of Study 1, but consisted only of a single safe block of 72 trials. This data was analysed immediately after completion of the calibration task in order to calculate the BP for each participant (i.e. the duration for which participants responded “short” or “long” equally often). This was set as the main stimulus duration for the fMRI experiment, in order to remove a potential confound of in Study 1: in that design, it was not clear whether the stimuli duration contrast indicated the neural correlates of how participants *perceived* time differently, or whether the neural effect was driven by the *actual presentation differences* between the stimuli. In order to minimize scanning time, participants performed this calibration task while anatomical brain images were acquired.

During the main task, each of the two runs consisted of six blocks (three safe and three threat blocks per run). Each block comprised 18 trials: the stimulus duration of 14 trials was set to the BP of each participant (calculated from the calibration task), the stimulus duration of two trials was 1.4s (the short “anchor” duration) and the stimulus duration of two trials was 2.6s (the long “anchor” duration). Thus, although in Study 1 and in the calibration task participants viewed six different durations (1.4, 1.64, 1.88, 2.12, 2.36, 2.6s), in the main task for Study 2 they viewed just three (1.4s, BP, 2.6s). Participants were told that in the main task the stimuli durations would be more difficult to tell apart compared to the calibration task. At the end of each block of the main task, a 20s rest period followed during which the screen went blank and participants were instructed to rest while the scan finished.

Participants received between zero and three shocks per block during the threat condition, according to a pre-determined schedule that resulted in participants receiving between 2-14 shocks in total, following a Gaussian distribution. The order of the shocks was random for each participant and occurred on different trials during the ITI. Each train of shocks consisted of 10 pulses delivered over 0.5 seconds.

Scanning was performed on a 3T Siemens Magnetom Prisma using a 64-channel head coil. The 1mm isotropic anatomical scan was a T1-weighted MPRAGE with the following parameters: TR=2.53s, TE=3.34ms, acquisition matrix=256 × 256, slice thickness 1mm, flip angle 7°. Functional (echo planar imaging: EPI) scans were acquired with a 2D sequence. Each volume consisted of 42 slices and was acquired in TR=2.94s, with ascending slice order. The angulation of the slice (T>C-30°), the phase-encoding direction and the compensating gradients (z-shimming) were optimized to minimize the dropout in regions near the orbito-frontal cortex and amygdalae (Weiskopf et al., 2006). Other parameters of the EPI were: voxel size 3mm × 3mm in-plane (slice thickness 2.5mm, 0.5mm slice separation); field of view 192mm×192mm; 12% over-sampling in the phase-encoded direction; bandwidth 2298 Hz/px; echo spacing 0.5ms, echo time 30ms; flip angle 90°. Fat saturation with an excitation of 130° was used prior to each excitation. At the end of each session, we acquired one fieldmap with identical parameters to the EPI scans. Heart rate and breathing were monitored using Spike2 software (http://ced.co.uk/products/spkovin).

### Behavioural data analysis

All data was processed in Matlab (v. R2015b), and statistical analysis was carried out in SPSS (v. 23).

#### Study 1

Trials on which participants did not make a response were excluded from the analysis. Repeated-measures analyses of variance (ANOVAs) were performed on the proportion of stimuli participants judged to be long (p(long)). The effects of threat (safe or threat-of-shock condition) and duration (six stimulus durations) were used as within-subject factors. Greenhouse-Geisser corrections were applied when violations of sphericity occurred. The BP was calculated for safe and threat conditions separately for each participant and the effect of threat was assessed using a paired-samples t-test.

#### Study 2

Trials on which participants did not make a response were excluded from the analysis. A paired-samples t-test was used to test the effect of threat on the proportion of long responses made at each participant’s BP (N=84 trials per condition). We did not analyse catch trials, i.e. the few trials (N=4 per block) whose durations were equal to the “short” (1.4s) and “long” (2.6s) anchors. However, we did use them to exclude one participant who pressed the same button on all trials, including both anchors, indicating that they did not understand/comply with the instructions.

### Functional neuroimaging data analysis

Study 1 was a pilot study which we used to generate ROIs for Study 2, and hence the data analysis for Study 1 was exploratory. In both studies, EPI data were analysed using Statistical Parametric Mapping (SPM12; Wellcome Trust Centre for Neuroimaging, London, www.fil.ion.uck.ac.uk/spm) in Matlab R2015b. After removing the first five volumes from each time series to allow for T1 equilibration, the remaining volumes were realigned to the sixth volume, normalized into standardized space (Montreal Neurological Institute (MNI) template), and smoothed using an 8 mm full width at half maximum Gaussian kernel. Following the realignment stage, all image sequences were checked for translations and rotations greater than 1.5 mm/1 degree – corrupted images were removed and replaced using interpolation. Following normalisation, images were manually checked for artefacts. Brain areas reported were defined using the Atlas of the Human Brain (Mai, 2016).

#### Study 1

Trials were modelled as events of zero duration. The regressors of interest were the onsets of the to-be-timed stimuli of all six durations (1.4, 1.64, 1.88, 2.12, 2.36, 2.6s) during safe and threat blocks (i.e. 12 regressors of interest). Regressors of no interest were the training stimuli indicating the anchor durations (presented before the beginning of each block), shocks, as well as the start screens of each block (indicating whether in that block participants were safe or under threat of shock). All these regressors were convolved with SPM’s canonical hemodynamic response function, time-locked to the onset of the corresponding event. We also included six movement regressors of no interest in all participants.

Using the general linear model, parameter estimate images were created for each regressor, and combined to create the primary contrasts at the subject level.

Second-level analyses were conducted using the standard summary statistics approach to random effects analysis. In this exploratory pilot study, we applied a lenient cluster-forming threshold of p<0.05 (uncorrected) cluster size>150, but did not use this to make inference. Instead we used the resulting clusters as ROIs in Study 2.

The fMRI contrasts were: 1) the effect of the threat compared to the safe condition; 2) the effect of stimulus duration, which was a linear contrast across the six stimulus durations (from short to long), collapsed across the safe and threat conditions; 3) the interaction of duration and threat.

#### Study 2

Due to changes in the experimental design we modelled the entire trial including stimulus presentation, stimulus response and ITI. Our four regressors of interest were threat trials (BP trials during the threat condition) and safe trials (BP trials during the safe condition), which were categorised as either “perceived long” (BP trials on which the participant responded with “long”) or “perceived short” (BP trials on which the participant responded with “short”). Regressors of no interest were the catch trials (i.e. trials whose duration was 1.4s and 2.6s) as well as missed trials (i.e. trials on which participant did not make a response), which were modelled separately for safe or threat blocks. Other regressors of no interest were training stimuli indicating the anchor durations (presented before the beginning of each block), shocks, as well as the start screens of each block (indicating whether participants will be safe or under threat of shock). All these regressors were convolved with SPM’s canonical hemodynamic response function time-locked to the onset of the corresponding event, and taking into account its duration (which varied slightly across participants due to variation in BPs).

We also included six movement regressors of no interest in all participants, alongside 12 regressors extracted from the pulse and respiratory rate, corresponding to a set of sine and cosine Fourier series components extending to the third harmonic (Glover et al., 2000) based on the Spike traces. There were also two regressors to model the variation in respiratory volume (Birn et al., 2006, 2008) and heart rate (Chang et al., 2009), also based on the Spike traces.

Using the general linear model, parameter estimate images were created for each regressor, and combined to create the primary contrasts at the subject level.

Second-level analyses were conducted using the standard summary statistics approach to random effects analysis. We applied a cluster-forming threshold of p<0.005 (uncorrected) and report small-volume corrected p-values for responses in our ROIs as defined in our pre-registration document (Sarigiannidis, 2019).

The fMRI contrasts were: 1) the effect of threat, i.e. all the BP trials under the threat-of-shock condition, compared to the safe condition; 2) perceived duration (“long” vs “short”, though the actual duration was identical), including all the BP trials, collapsed across the threat and safe conditions; 3) the interaction between perceived duration and threat. We additionally examined the overlap between (1) and (2) using conjunction analysis.

#### Regions of Interest

In Study 1 we detected significant threat-induced activation in the anterior cingulate cortex, and since this area was also activated during threat-of-shock conditions in our group’s previous studies (Robinson et al., 2014) we used it as a pre-registered ROI. However, due to a mix-up the co-ordinates identified as the anterior cingulate cortex in the pre-registration document actually refer to the left caudate because both peaks fell within the same large cluster. In the interests of full transparency, we therefore used both ROIs for the threat contrast using a 10-mm sphere for both the caudate (MNI coordinates [x=-18, y=11, z=26]) and the anterior cingulate cortex (MNI coordinates [x=0, y=-4, z=50]; prediction 2).

Previous meta-analysis on time perception studies reported strong activation in the pre-SMA (Wiener et al., 2010), and thus for contrast (1) we defined an additional ROI defined as a 10-mm sphere ROI on that area (again this was pre-registered: Talairach coordinates [x=0, y=0, z=56], taken from Wiener et al. (2010), converted to MNI coordinates [x=-1, y=-4, z=62]; prediction 3).

## Results

### Study 1

#### Behavioural results inside the scanner

Across blocks, participants reported being significantly more anxious in the threat compared to the safe condition (t(12)=5.40, p<0.001, d=1.56). There was a significant effect of stimulus duration (F(2.46, 29.58)=211.51, p<0.001, η_p_^2^=.946; see Figure 2). As expected, the longer the stimulus duration, the more likely it was to be classified as “long”. The effect of threat was non-significant (F(1, 12)=2.57, p=0.135, η_p_^2^=.177) and neither was the threat-by-duration interaction (F(5, 60)=0.27, p=0.926, η_p_^2^=.022). The BP was not significantly different between the threat and safe conditions (t(12)=1.36, p=0.194, d=0.37), although the direction of the effect was consistent with our previous studies (a rightward shift in the psychometric function, consistent with temporal underestimation during threat). This non-significant result is expected since the study is underpowered, and this was not the purpose of the study.

**Figure 2:**
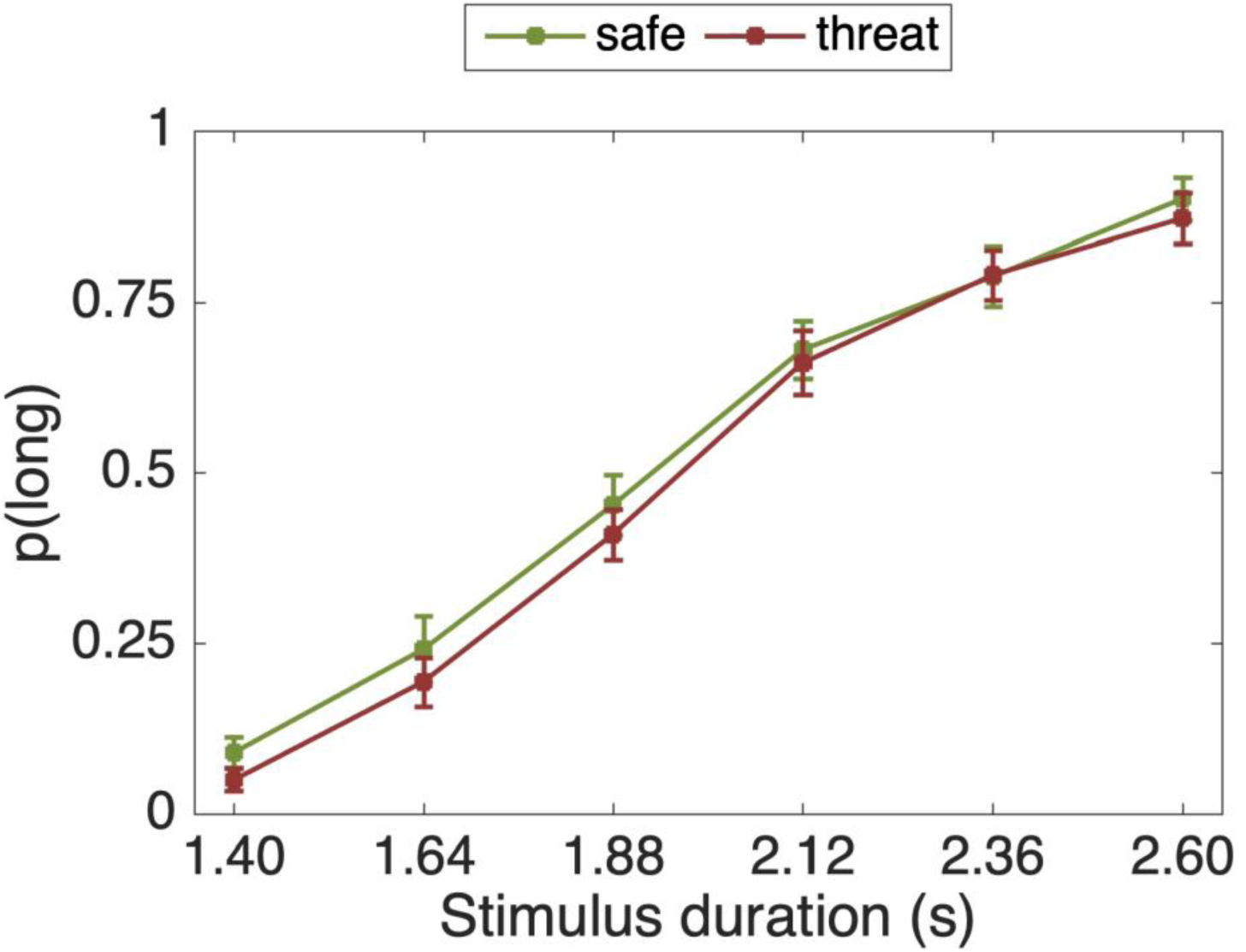
Proportion of stimuli rated “long” as a function of the actual presentation length and threat condition. Error bars are standard errors of the mean (SEM).

#### Neural effect of threat

No voxels survived correction for multiple comparisons in the threat contrast, the duration contrast or the interaction. Therefore, the below analyses were conducted using an exploratory threshold of P<0.05 (uncorrected), and used to generate hypotheses for Study 2. Only large clusters (k>150) are reported (see Table S1)

##### Threat>safe

This analysis examined the effect of the threat-of-shock vs the safe condition and revealed activations in two large clusters. One cluster was in the parietal cortex, consisting mainly of white matter and therefore was not considered further. The peak activation in the other cluster was in the left caudate ([x=-18, y=11, z=26) while in the same cluster, there was also a sub-peak in anterior cingulate cortex ([x=0, y=-4, z=50]); see Figure S1A. Both of these areas were used as ROIs for the threat contrast of Study 2.

##### Safe>threat

This analysis examined the effect of the safe vs the threat-of-shock condition and revealed activations in one large cluster, with a peak in the right supramarginal gyrus.

#### Neural effect of stimulus duration

##### Long>short

This analysis examined the effect of stimulus duration implementing a linear contrast (−2.5, - 1.5, −0.5, 0.5, 1.5, 2,5 for the six stimulus durations: 1.4, 1.64, 1.88, 2.12, 2.36, 2.6s) and revealed activation in one large cluster, with a peak activation in the visual cortex (right superior occipital gyrus; see Figure S1B).

##### Short>long

This analysis examined the effect of stimulus duration implementing the inverse of the above contrast which revealed activation in one large cluster, with a peak activation in the left superior cerebellar peduncle.

#### Neural effect of threat × stimulus duration interaction

The analysis examined the interaction of the above effects, revealing activation in one large cluster, with a peak activation in the precuneus. There was also activation in the frontal areas, specifically in the left middle frontal gyrus (see Figure S1C). The inverse contrast revealed activation in one large cluster, with a peak activation in the precentral gyrus.

### Study 2

#### Behavioural results inside the scanner

Participants reported being significantly more anxious in the threat compared to the safe condition (t(28)=11.28, p<0.001, d=2.09). As hypothesised, on BP trials participants responded “short” significantly more often in the threat compared to the safe condition (t(28)=2.39, p=0.024, d=0.44; Figure 3).

**Figure 3:**
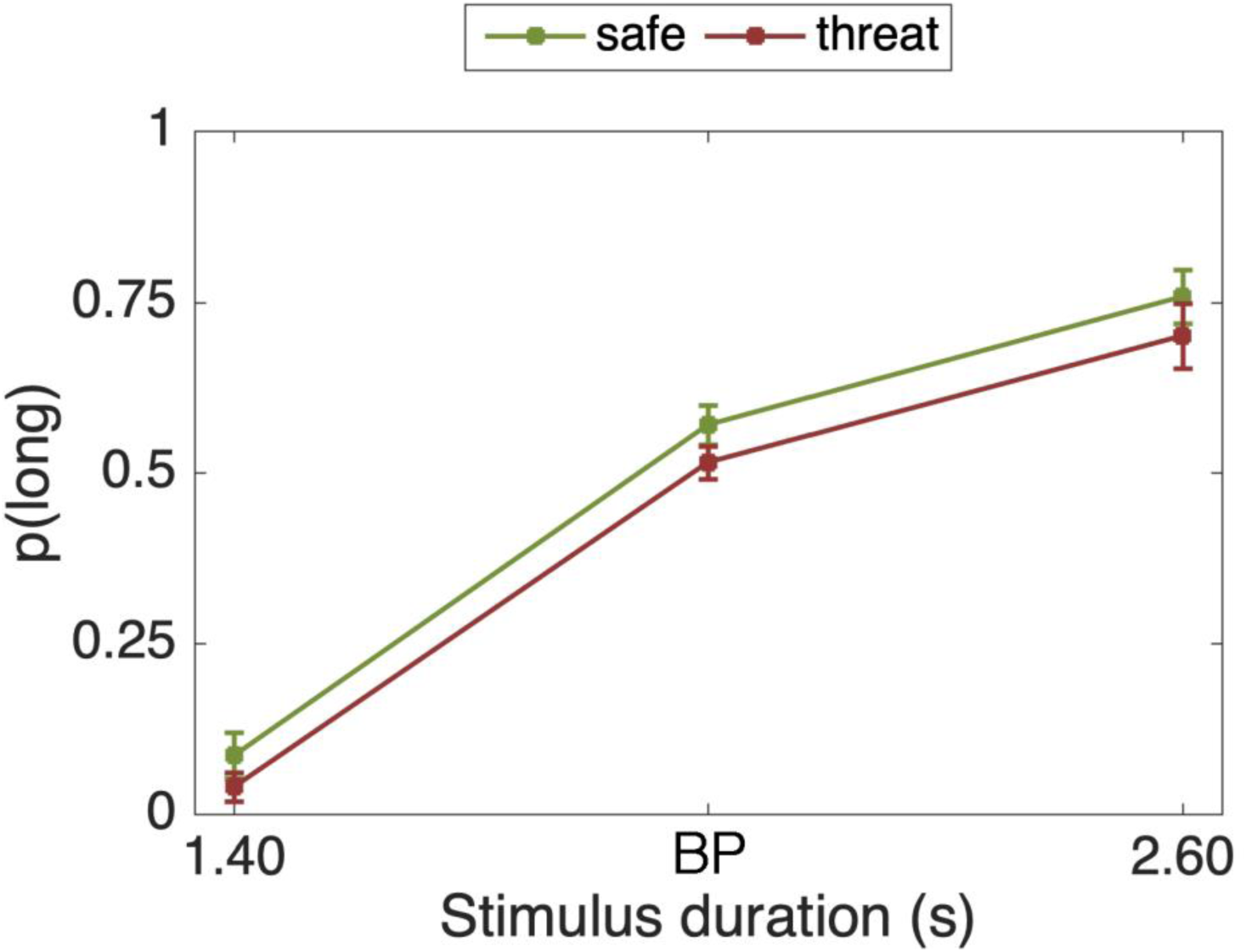
Proportion of p(long) responses was significantly lower in the threat condition for the individually tailored bisection point (BP) duration, suggesting temporal underestimation during anxiety. Error bars are standard errors of the mean (SEM). The 1.4 and 2.6s durations were not used in any analyses and are only plotted here for completeness.

#### Neural effect of threat

##### Threat>safe

This analysis examined the effect of the threat-of-shock vs the safe condition. There was significant (whole-brain voxel-level FWE corrected) activation in a large cluster (see Figure 4 and Table 2), including peaks in the subgenual anterior cingulate cortex (bilateral), thalamus (bilateral), claustrum (left only), caudate (left only) and anterior insula (bilateral). Other peaks in this cluster that did not survive voxel-level correction were an insula/orbitofrontal cortex area (right only), the lateral septal area (right only) and the putamen (right only). Three more significant clusters revealed activations in the left cerebellum and the parietal operculum (left and right).

**Table 1:** Summary of studies conducted. Note that the studies were completed at different sites in different countries.

|  | Sample size | Stimulus duration | Shock type | BDI | STAI |
| --- | --- | --- | --- | --- | --- |
| Study 1:<br>NIH | 13 | 1.4-2.6s (1.4, 1.64, 1.88, 2.12, 2.36, 2.6s) | single pulse | n/a | n/a |
| Study 2:<br>UCL | 29 | BP (78% trials of interest),<br>1.4s & 2.6s (22% catch trials) | train of shocks<br>(0.5s) | 5.06<br>(5.53) | 35.72<br>(12.48) |

**Table 2:**
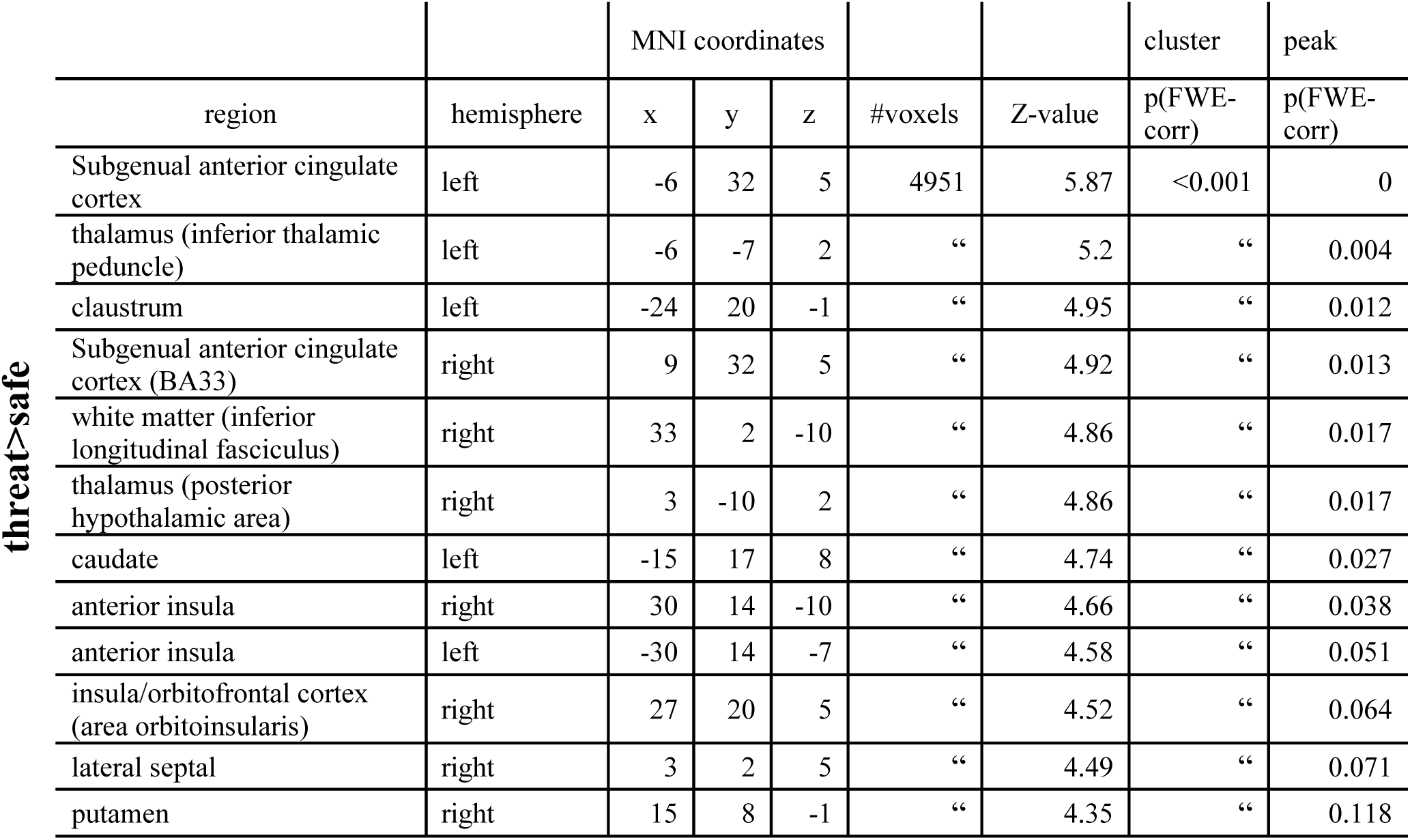
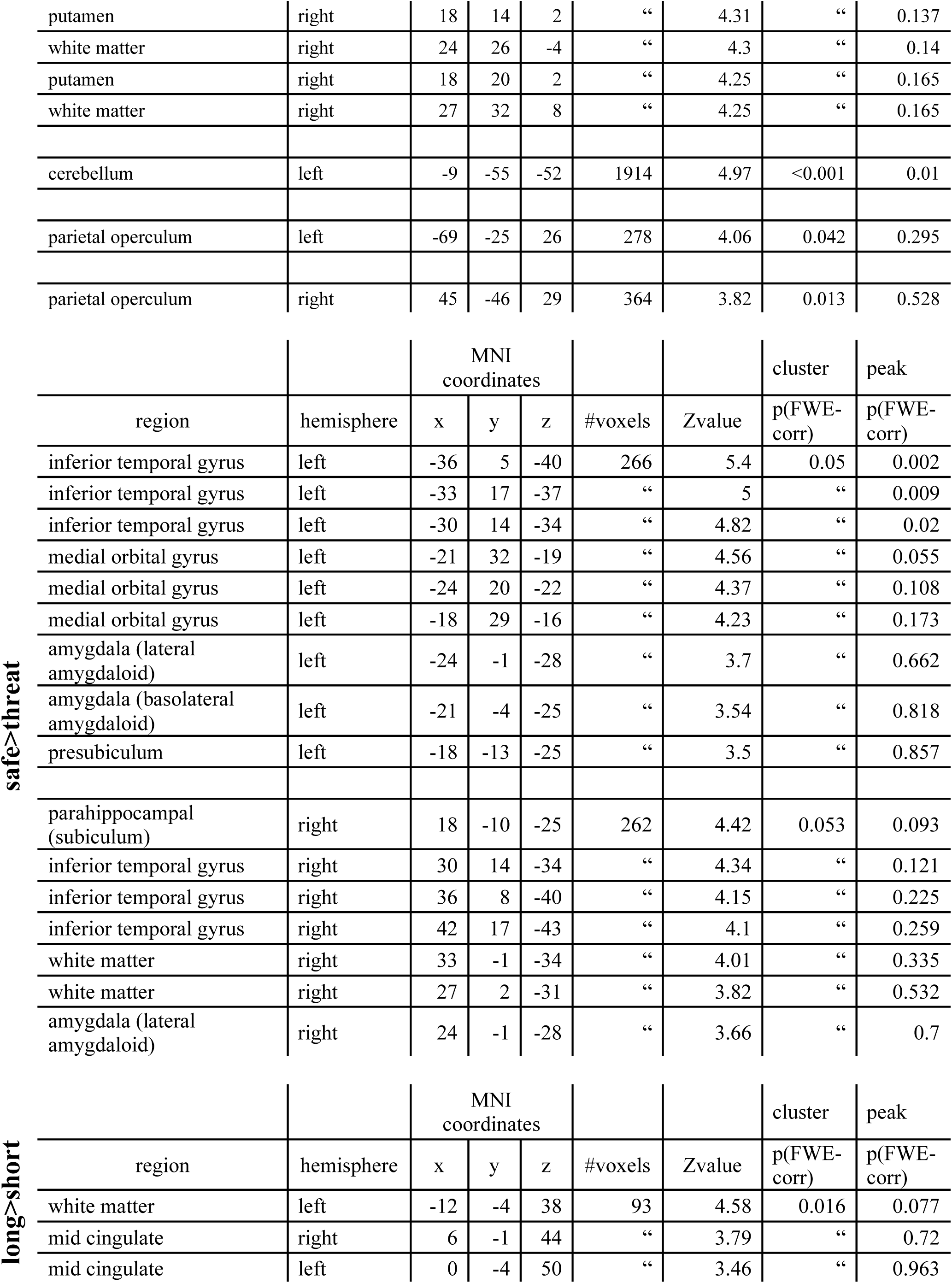
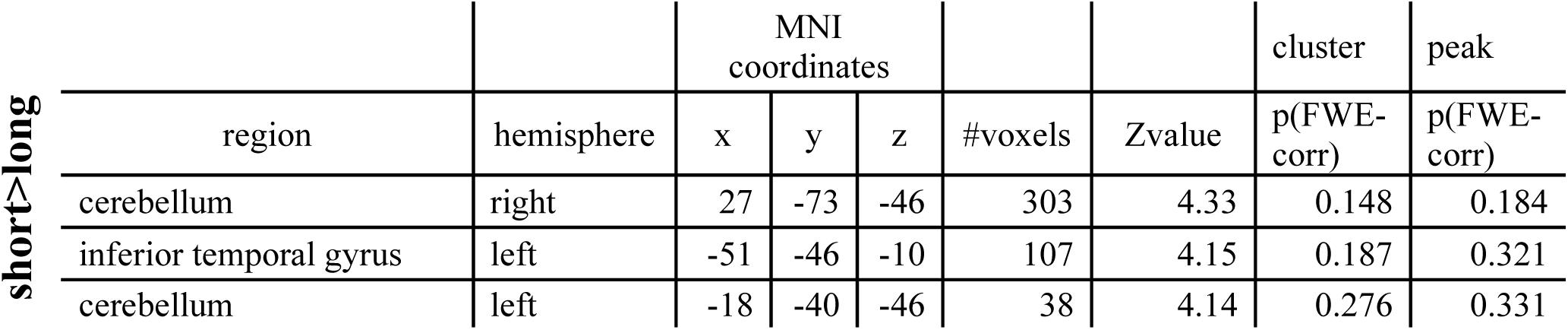
fMRI activations for [threat>safe,] [safe>threat], [perceived long>perceived short], and [perceived short> perceived long] contrasts (cluster forming threshold: p<0.005, uncorrected).

**Figure 4:**
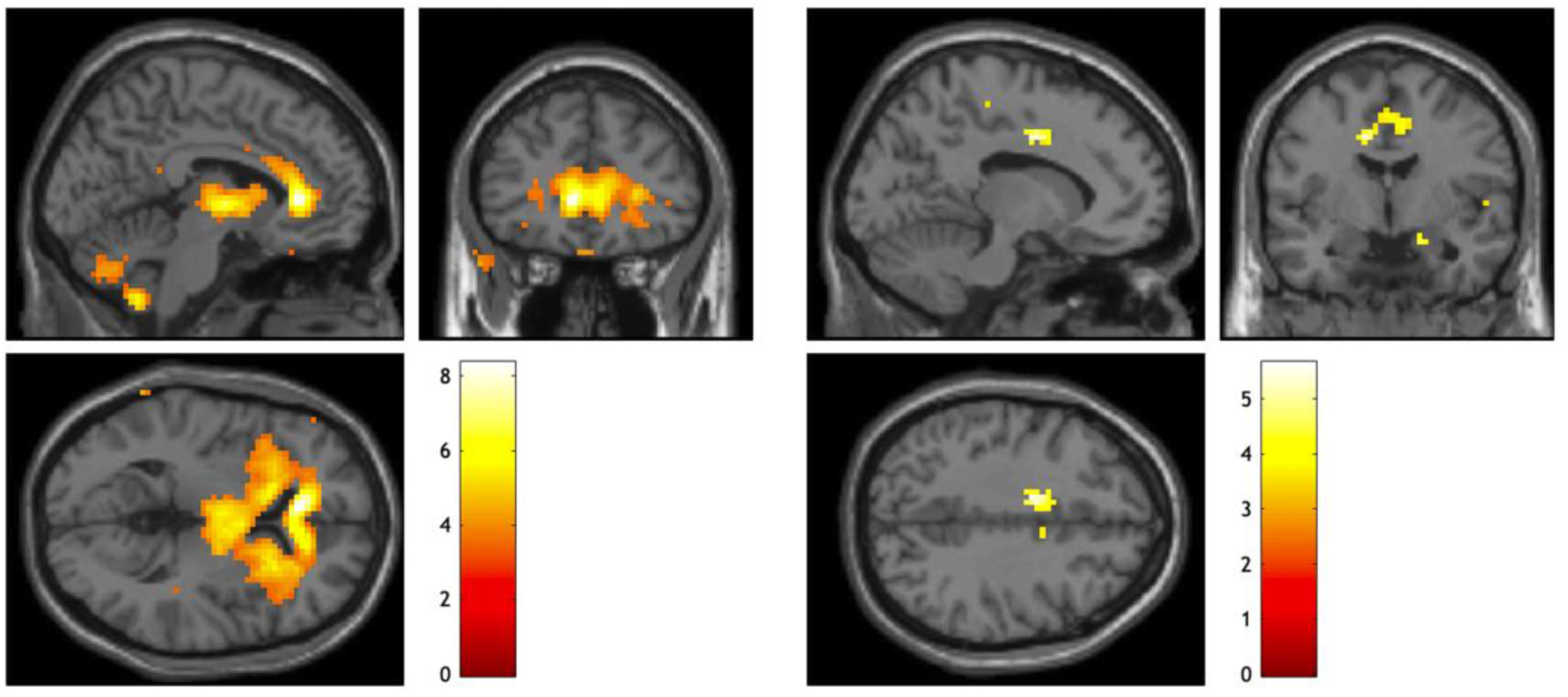
Activation for the threat>safe contrast (left) and the perceived long> perceived short trials (right). Cluster forming threshold p<0.005 (uncorrected). The colour bar represents t-values.

In our pre-registration document, we defined the caudate and the anterior cingulate cortex as ROIs, since they were both activated under threat-of-shock in Study 1. When small volume correction was applied using a 10-mm sphere ROI around the peak of the caudate cluster identified in Study 1 (MNI coordinates [x=-18, y=11, z=26]), a peak survived FWE voxel-level correction for multiple comparisons ([x=-15, y=20, z=23], Z=3.21, k=37, p<0.05). The ROI around the peak of the anterior cingulate cortex cluster from Study 1(MNI coordinates [x=0, y=-4, z=50]), also revealed a peak surviving FWE voxel-level correction for multiple comparisons ([x=3, y=-4, z=41], Z=2.90, k=11, p<0.05).

Additionally, to explore brain-behaviour correlations we performed an exploratory analysis in which the effect of threat on behavioural responses (p(Long)_threat_- p(Long)_safe_) was entered as a covariate into the [threat>safe] contrast. We expected that the neural effect of threat>safe would be larger in participants who showed greater temporal underestimation during threat. However, no activations survived correction for multiple comparisons. Equally, no activations survived correction in the inverse contrast.

##### Safe>threat

This analysis examined the effect of the safe vs the threat-of-shock condition. There was significant (whole-brain voxel-level FWE corrected) activation in two clusters with bilateral peaks in the inferior temporal gyrus, in the left medial orbital gyrus (see Figure 4 and Table 2), and in a right parahippocampal area (subiculum). Both clusters extended into the amygdalae, although the peaks there did not survive voxel-level correction.

#### Neural effect of perceived duration

##### Perceived long> perceived short trials

This analysis examined the effect of the perceived long vs. short trials, i.e. trials in which participants judged the to-be-timed stimulus (which was actually always the same duration at their own BP) as “long” compared to “short”. There was significant activation in a single cluster with bilateral peaks in mid-cingulate cortex (see Figure 4 and Table 2).

In our pre-registration document we defined the supplementary motor area as an ROI since it has been reliably implicated in time perception studies. When small volume correction was applied using a 10-mm sphere ROI around the meta-analytic peak identified by Wiener et al (2010: MNI coordinates [x=-1, y=-4, z=62]) we identified a peak which survived FWE voxel-level correction for multiple comparisons ([x=0, y=-4, z=53], Z=3.14, k=11, p<0.05).

Additionally, in an exploratory analysis, we investigated whether activation in the mid-cingulate cortex peak voxel was associated with the degree of temporal underestimation during threat, but the correlation was non-significant (r(29)=-.28, p=.884).

##### Perceived short> perceived long trials

This analysis examined the effect of the perceived short vs. long trials, i.e. trials in which participants judged the to-be-timed stimuli as “short”, compared to when they judged them as “long”. No clusters survived correction at either the peak or voxel level (Table 2).

#### Neural effect of threat × perceived duration

This analysis examined the interaction between the effect of threat and that of perceived duration. No clusters survived correction in either this or the inverse contrast.

#### Overlap analyses

The results of Study 2 suggest a degree of overlap in the activations identified in the [threat>safe] and [perceived long>perceived short] contrasts. We formally tested this overlap by creating a mask for each contrast (thresholded at t>1.7, corresponding to p<0.05 uncorrected), using this to perform small volume correction on the other contrast (see Figure 5), using a cluster-forming threshold of p<0.005 (uncorrected).

**Figure 5:**
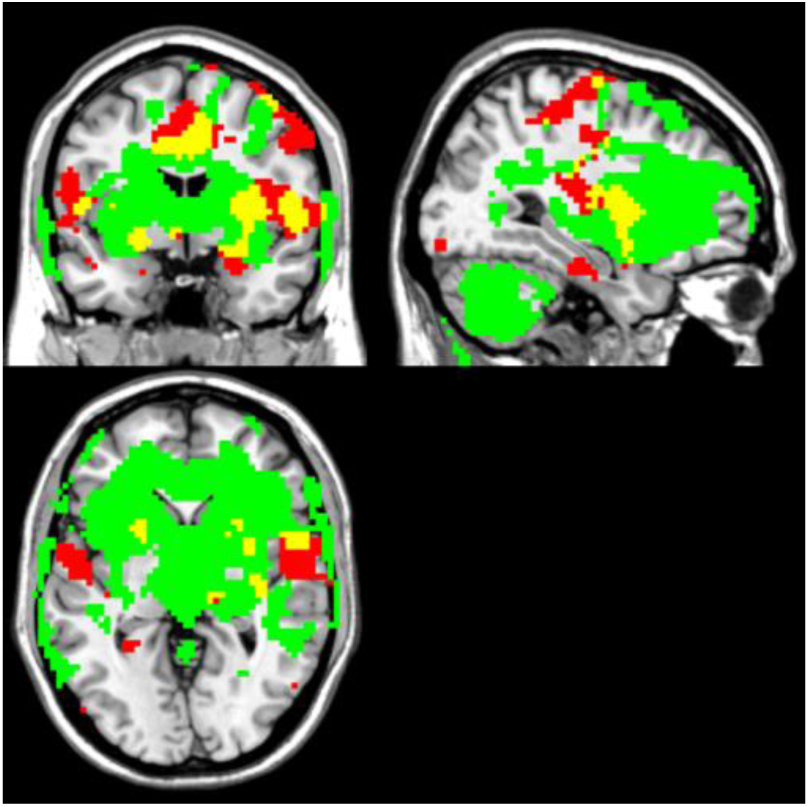
Overlapping activations of [threat>safe] (in green) and [perceived long>perceived short] contrasts (in red). Overlapping regions (in yellow) include the insula ([x=-27, y=8, z=-7]), putamen ([x=30, y=2, z=-7]) and mid-cingulate ([x=-12, y=2, z=38]). Figure generated by creating masks from the [threat>short and [perceived long>perceived short] contrasts, both thresholded at p<0.05 (uncorrected) for display purposes Note: activation of other brain areas has been omitted for clarity.

Applying the [perceived long>perceived short] mask to the [threat>safe] contrast revealed significant overlap surviving FWE voxel-level correction for multiple comparisons in right insula ([x=30, y=2, z=-7], Z=4.35, k=57, p<0.05) and left putamen ([x=-27, y=8, z=-7], Z=4.03, k=12, p<0.05).

Applying the [threat>safe] mask to the [perceived long>perceived short] contrast revealed overlap in a mid-cingulate cortex area which narrowly missed FWE voxel-level correction for multiple comparisons ([x=-12, y=2, z=38], Z=4.29, k=15, p=0.068).

To test our hypothesis that anxiety may “overload” regions processing time perception, we performed a 2-by-2 (threat-by-perceived duration) ANOVA on average activation across the above ROIs (Figure 6). However, there was no significant interaction between threat and perceived duration for either the insula (F(1,28)=.049, p=.827) or the mid-cingulate (F(1,28)=.075, p =.786). We report only these interactions here, and not the main effects, to avoid circularity in the analysis. The main effects of threat and perceived durations have already been reported in previous sections.

**Figure 6:**
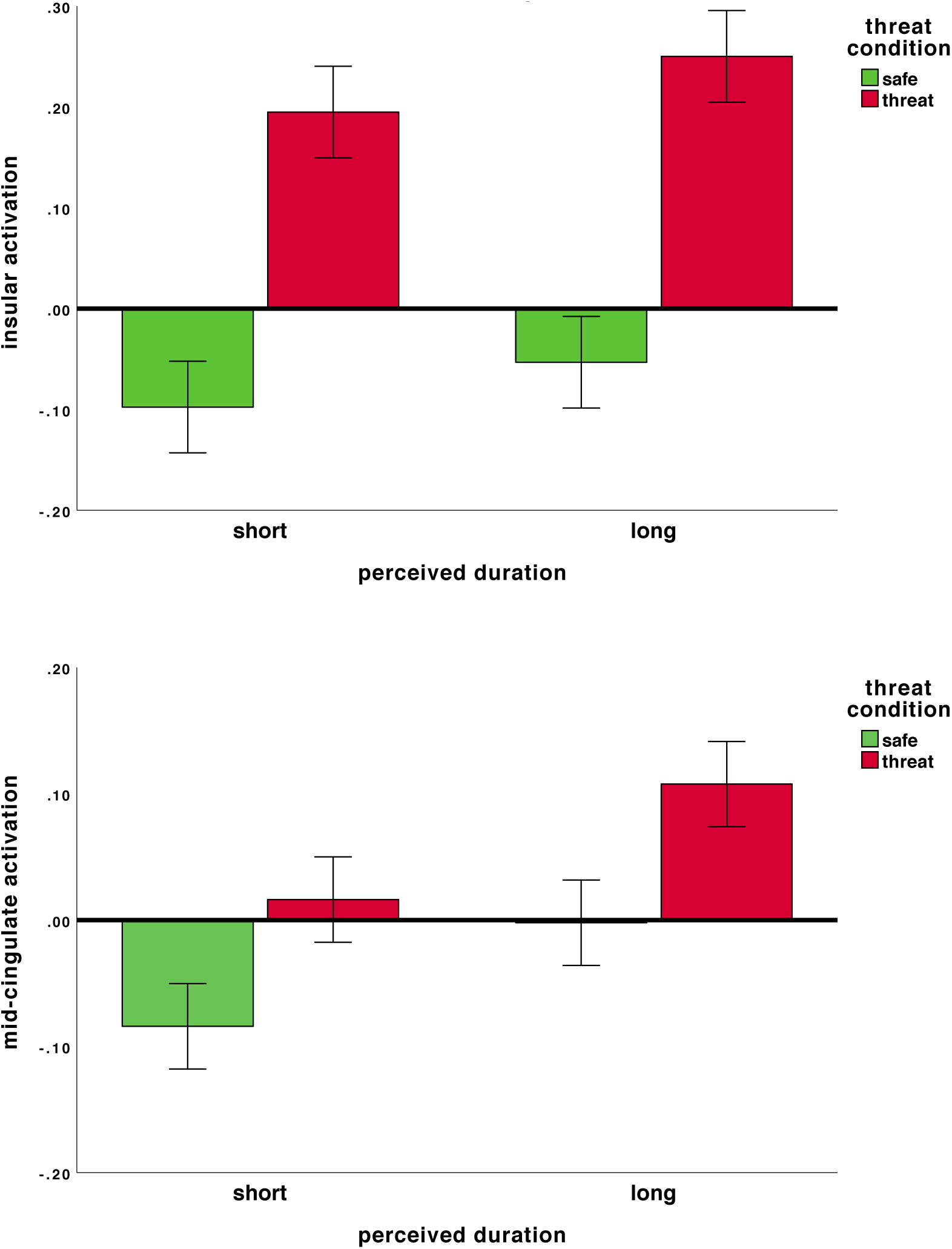
Averaged activation across the ROI for each condition in the right insula (top panel) and the mid-cingulate (bottom panel). Errors bars are standard errors.

One possibility is that individuals whose task responses were affected by anxiety (i.e. underestimated time), would show an interaction between the insula and the mid-cingulate, due to neural overloading. At the same time, we would not expect an interaction on individuals who did not underestimate time under threat. Hence, we extracted the activation of the insula and mid-cingulate areas from the interaction contrast, and correlated it with the degree of temporal underestimation during threat. No significant correlations were observed, either between the effect of threat on pLong and the right insula from the interaction contrast (r(29)=.16, p=.51), or between the effect of threat on pLong and the mid-cingulate from the interaction contrast (r(29)=.07, p=.72)

## Discussion

We explored the neural correlates of how anxiety alters cold cognition by combining a threat-of-shock manipulation and a temporal bisection task. We replicated and extended our previous behavioural findings (Sarigiannidis, Grillon, et al., 2020) in the scanner, showing that participants underestimated the duration of a single temporal interval (corresponding to their BP) when anxious (prediction 1). We further found that induced anxiety activates the anterior cingulate cortex and caudate (prediction 2), as well as the insula, consistent with previous studies (Balderston et al., 2017; McMenamin et al., 2014; Meyer et al., 2019; Robinson et al., 2014, 2019; Torrisi et al., 2016, 2018). Additionally, although the perception of longer temporal intervals was associated with activation in the pre-SMA (prediction 3) consistent with previous studies (Schwartze et al., 2012; Wiener et al., 2010) this ROI analysis did not survive in the whole brain analysis. Instead, a mid-cingulate area was more robustly activated when participants perceived the temporal intervals as long. Finally, in-line with an “overloading” hypothesis, activations in the threat and perceived duration contrasts overlapped in the insula and mid-cingulate area (but not in the pre-SMA; prediction 4).

### Neural correlates of anxiety induced by threat of shock

The pattern of anxiety-specific neural activations (threat>safe) was largely consistent with previous studies. Specifically, the whole brain analysis in Study 2 revealed a large cluster of activation in the ACC, with a peak in the sgACC. This widespread anxiety-related activation included a caudate and an ACC area identified in Study 1, which we confirmed using small-volume correction, confirming prediction 2.

Since our participants received more electrical shocks than in previous threat-of-shock studies (e.g. Balderston et al., 2017; Robinson et al., 2014; Torrisi et al., 2016), it is possible that the sgACC activation is due to processing painful stimuli (for a review see Palomero-Gallagher et al., 2015). However, we did regress out the effect of shocks, hence activation in this region is most likely due to shock anticipation. This is consistent with sgACC activation being considered central in sustained anticipatory responses in both primates and humans (Robinson et al., 2014; Rudebeck et al., 2014; Wallis et al., 2017).

At the same time, the right insula and the right caudate were activated across Study 1 and 2. These areas have previously been implicated in induced anxiety (insula: Balderston et al., 2017; McMenamin et al., 2014; Meyer, Padmala, & Pessoa, 2019; caudate: McMenamin et al., 2014; Torrisi et al., 2016). Insula is often co-activated with the ACC (Palomero-Gallagher et al., 2015) and is considered to be part of a putative “anxious anticipation” network (McMenamin et al., 2014). Although the caudate is less consistently implicated in threat-of-shock studies, a previous study has similarly found activation in the right caudate (Torrisi et al., 2016). Considering that the caudate is considered key area in pathological anxiety, constituting targets for deep brain stimulation of disorders involving anxiety (i.e. obsessive compulsive disorder; Alonso et al; 2015) future studies could further explore how it relates to anticipatory anxiety. Taken together, these results highlight important roles of sgACC, the insula and the caudate in anticipatory anxiety.

### Neural correlates of temporal perception

Our results suggest that a mid-cingulate area was more active when participants perceived a stimulus as “long” than when they perceived the exact same stimulus as “short”. It is possible that this mid-cingulate area is involved in monitoring stimulus duration, where increased activation reflects more efficient processing (i.e. fewer temporal pulses were lost). The cingulate cortex has previously been implicated in time perception (for a recent meta-analysis see Nani et al., 2019), but less consistently than other brain areas such as the pre-SMA. This apparent discrepancy might be attributed to the different experimental tasks used.

Specifically, previous fMRI studies on perceptual timing have mainly employed comparative temporal discrimination tasks, in which participants judge which of the two consecutively presented temporal intervals was longer. Thus, in these studies the neural signal may represent general perceptual timing, including processes such as keeping track of different temporal intervals and working memory; since durations have to be kept in mind to allow comparisons on each trial. In our task, participants viewed the exact same temporal interval which they compared with temporal durations they had consolidated (i.e. the anchor durations); hence the neural signal reflects differences in perception free of working memory confounds or any other confounds related to stimulus duration.

Nevertheless, the pre-SMA has been implicated across different temporal cognition tasks, from motor (e.g. finger tapping) to perceptual and is considered a key area in timing (Nani et al., 2019; Schwartze et al., 2012; Wiener et al., 2010). In our study, our pre-registered ROI analysis did confirm activation in the pre-SMA (prediction 3). However, it did not survive correction in the whole-brain analysis, which suggests that other regions may be more important in our study. This raises questions about precisely what role the pre-SMA plays in keeping track of time. It is also possible that the pre-SMA participates in some general aspect of temporal processing, such as using strategies to count interval durations, which might explain why it is so ubiquitously activated across so many different temporal cognition tasks (Nani et al., 2019).

Finally, we found overlapping mid-cingulate cortex activation in the threat and perceived duration contrasts. This convergence raises the possibility that mid-cingulate cortex might be implicated in emotion-related alterations in temporal perception, in-line with the hypothesised role of this region in mediating cognitive affective and behavioural responses to anxiety (Grupe & Nitschke, 2013).

### Neurocognitive mechanisms of temporal underestimation under anxiety

We previously hypothesised that the effect of anxiety on temporal cognition was due to dual task interference: anxiety may occupy limited neurocognitive resources, thus altering performance in the temporal estimation task (Sarigiannidis, Grillon, et al., 2020; Sarigiannidis, Kirk, et al., 2020). Specifically, in this study we considered the threat-of-shock condition to represent a dual-task scenario, since participants are performing the temporal task whilst also “processing” anxiety, and the safe condition to be single-task, since participants are only performing the temporal task.

We found preliminary evidence that insula and a mid-cingulate area were activated both during the threat and temporal contrasts. It is thus possible that anxiety-related insula activity (Baur et al., 2013; Bijsterbosch et al., 2015; Simmons et al., 2006) interfered with the mid-cingulate cortex (an area associated with time perception Nani et al., 2019), leading it to less efficiently accumulate temporal information (e.g. losing temporal pulses) and thus resulting in the temporal underestimation we observed. However, this hypothesis is not completely supported by our data, considering that we did not find a significant threat-by-perceived duration interaction either at the whole-brain level, or when specifically examining the insula or mid-cingulate. In fact one might expect no interaction between the insula and mid-cingulate activation in the interaction contrast in participants without threat-induced time underestimation; but an interaction (due to overloading) in those who did underestimate time under threat. However, our data did not support this either: we did not detect any correlation across participants between the underestimation of time during threat and either insula or mid-cingulate activation in the interaction contrast. Future work could explore this hypothesis including more participants to provide adequate power to detect individual differences (rather than just within-subject differences) and by employing a similar task with a fully factorial design, additionally incorporating a control/passive task. This might provide a more complete picture of how anxiety and cognition interact at the neural level.

Previous studies have suggested that the dorsolateral prefrontal cortex is implicated in the cognitive alterations due to trait (Bishop, 2009) and induced anxiety (Balderston et al., 2017; Torrisi et al., 2016). However, we did not find such activation here. This discrepancy can be explained considering that the tasks utilised in the aforementioned studies involved cognitively demanding, fast-paced tasks which were more likely to activate prefrontal areas (Höller-Wallscheid et al., 2017) than our simple task. Taken together our results tentatively support the idea of anxiety altering cognition similarly to dual-task situations, but in our data there was no evidence that this was mediated by dorsolateral prefrontal activation.

### Limitations

Study 1 was a pilot study with a different design to Study 2. In Study 1 participants had to judge the duration of six different temporal intervals. This version of the task confounded duration and time perception, such that effects might simply be driven by the duration of the stimuli. In other worlds, it is not clear whether the stimulus duration contrast indicates the neural correlates of how participants *perceived* time differently, or whether the neural effect was driven by the longer presentation times. There is evidence consistent with the latter explanation, since in the contrast of increasing stimulus duration in Study 1 we identified activation in the visual cortex, which could be explained by a purely sensory account.

This confound was eliminated in Study 2 where participants had to judge the duration of a fixed duration stimulus. This was a stimulus duration that tailored to participants responding equally frequently to “short” or “long” (calculated from a calibration task similar to Study 1) and thus any neural differences found in this contrast corresponds to how participants *perceived* time, free of any confounds of the *actual* duration of the stimuli (which was identical throughout). In Study 2 we did not identify any visual cortex activation, consistent with the above explanation.

We also failed to find any correlations between a) the behavioural effect of threat and the neural effect of threat>safe and b) the behavioural effect of threat and neural activation in the mid-cingulate area that was active during both the threat>safe and the perceived long>perceived short contrasts (exploratory analyses). This may be due to low statistical power, as our study was not optimised to examine individual differences. Indeed, in order to detect a correlation of .30 with 0.80 power we would need 82 participants, which was beyond the scope of the current within-subjects design.

It should also be noted that our fMRI paradigm does not allow us to dissociate between perception of the stimulus and the response, thus all interpretations should be made with this in mind. A future study could further delineate this by introducing a jittered delay between stimuli perception and task response in the design of the task. This (longer) design would enable the investigation of how anxiety affects time perception at the stage of perception and/or decision and which neural circuits are involved at each stage.

Finally, we tested healthy individuals under an anxiety manipulation. Whether our results would generalise to pathological anxiety remains an open question.

### Conclusion

We replicated previous findings of temporal underestimation in anxiety, and found activation in brain areas previously associated with threat-of-shock-induced anxiety (sgACC, insula, caudate) and time perception (mid-cingulate cortex). Despite previous studies suggesting a key role of pre-SMA in temporal perception, the mid-cingulate cortex was more robustly activated in our study. We suggest a potential role of the mid-cingulate cortex in mediating emotion-related alterations in temporal perception. Interestingly, we found evidence of overlap between activations elicited by time perception and anxiety (insula and mid-cingulate cortex), which is consistent with the hypothesis that anxiety may influence cognition by further loading already-in-use resources.

## Author contributions

I. Sarigiannidis developed the study concept, who designed the study with input from O. J. Robinson and J. P. Roiser. Testing and data collection were performed by I. Sarigiannidis under the supervision of O. J. Robinson and J. P. Roiser at UCL (Study 2), and Christian Grillon and Monique Ernst at NIH (Study 1). K. Kieslich assisted with testing and data collection of Study 2. Data analysis and interpretation was carried out by I. Sarigiannidis, O. J. Robinson and J. P. Roiser. I. Sarigiannidis drafted the manuscript with critical revisions from O. J. Robinson and J. P. Roiser. Christian Grillon and Monique Ernst provided useful comments to this draft. All authors approved the final version of the manuscript for submission.

## Acknowledgements

We would like to thank Joe Devlin, Steve Fleming and Karl Friston for useful comments regarding the design of Study 2.

