## Supplementary Material for "Anxiety makes time pass quicker: neural correlates"

### Study 1 Analysis

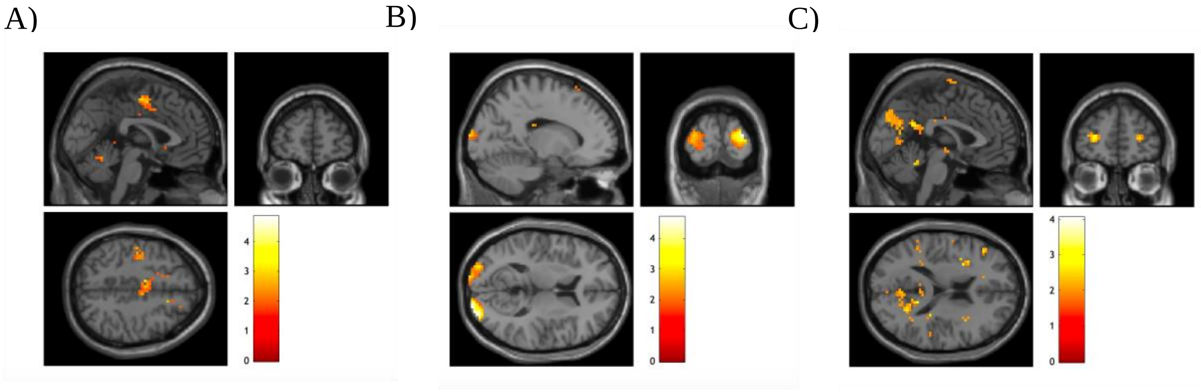


Figure S1: Uncorrected, exploratory BOLD activation for each contrast in Study 1: A) threat>safe; B) linear effect of lengthening temporal interval; C) positive interaction of A and B. A threshold of P<0.05 (uncorrected) was used, colour bars indicate t-values.

Table S1: Activations for the Study 1 contrasts, exploratory cluster forming threshold p<0.05 (uncorrected)

| | | | MNI coordinates | | | | | cluster | peak |
|---|---|---|---|---|---|---|---|---|---|
| contrast | region | hemisphere | x | y | z | #voxels | Zvalue | p(FWE-corr) | p(FWE-corr) |
| threat>safe | white matter | n/a | 0 | 27 | -7 | 390 | 3.51 | 0.136 | 1 |
| | caudate | left | -18 | 11 | 26 | 240 | 3.26 | 0.626 | 1 |
| | anterior cingulate | n/a | 0 | -4 | 50 | “ | 2.51 | “ | 1 |
| safe>threat | supramarginal gyrus | right | 36 | -43 | 35 | 272 | 3.77 | 0.471 | 0.997 |
| long>short | occipital gyri | right | 24 | -100 | 8 | 247 | 3.47 | 0.989 | 0.992 |
| short>long | superior cerebellar peduncle | left | -3 | -25 | -13 | 24626 | 4.56 | <0.001 | 0.154 |
| interaction | precuneus | left | -9 | -67 | 44 | 446 | 2.63 | 0.491 | 1 |
| | frontomarginal gyrus | left | -24 | 46 | 2 | 66 | 3.15 | 1 | 1 |

| | | | | | | | | | |
|---|---|---|---|---|---|---|---|---|---|
| interaction (inverse) | precentral gyrus | right | 59 | -2 | -23 | 182 | 2.88 | 0.997 | 1 |

**Overlap between Study 1 & 2**

A whole-brain mask (thresholded at p<0.05 uncorrected) was created from each contrast of Study 1 (using the ImCalc function in SPM) and was applied to the corresponding contrast for Study 2, using P<0.005 as the cluster-forming threshold and a 10-voxel cluster size. Statistical inference was based on the peak level statistics, with voxel-level small volume correction applied across the mask.

When applying a mask generated from the threat > safe contrast in Study 1 to the equivalent contrast in Study 2 there was significant (voxel-level FWE corrected) activation in the right insula and the right caudate. There was no significant overlap between Studies 1 & 2 for the duration or interaction contrasts.

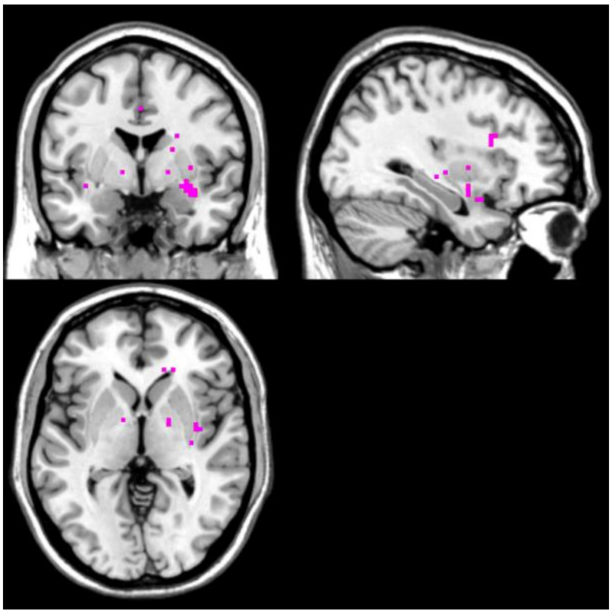

Figure S2: BOLD co-activations for threat>safe contrasts for Study 1 and Study 2. Figure generated by creating masks from the threat>short contrast for Studies 1 & 2, thresholded at t>1.78 and t>2.76 respectively. Left panel: insula, right panel: caudate

Table S2: fMRI activation overlap between Study 1 & 2 for threat>safe.

| | | MNI coordinates | | | | | cluster | peak |
|---|---|---|---|---|---|---|---|---|
| region | hemisphere | x | y | z | #voxels | Zvalue | p(FWE-corr) | p(FWE-corr) |
| insula | right | 33 | -1 | -10 | 11 | 4.51 | 0.754 | 0.003 |
| white matter | right | 18 | 38 | 5 | 1 | 4.25 | 0.911 | 0.009 |
| lateral caudate nucleus | right | 18 | 20 | 17 | 49 | 4.10 | 0.354 | 0.017 |
| white matter | right | 27 | 20 | 23 | “ | 4.04 | “ | 0.022 |
